# Symmetry breaking and *de-novo* axis formation in *hydra* spheroids: the microtubule cytoskeleton as a pivotal element

**DOI:** 10.1101/2020.01.14.906115

**Authors:** Heike Sander, Aravind Pasula, Mathias Sander, Varun Giri, Emmanuel Terriac, Franziska Lautenschlaeger, Albrecht Ott

## Abstract

The establishment of polarity in cells and tissues is one of the first steps in multicellular development. The ‘eternal embryo’ *hydra* can completely regenerate from a disorganized cell cluster or a small fragment of tissue of about 10, 000 cells. During regeneration, the cells first form a hollow cell spheroid, which then undergoes *de-novo* symmetry breaking to irreversibly polarize. Here, we address the symmetry-related shape changes. Prior to axis establishment, the spheroid of regenerating cells presents inflation oscillations on several timescales that are isotropic in space. There are transient periods of fluctuations in defined arbitrary directions, until these undergo a clearly identified, irreversible transition to directed fluctuations along the future main axis of the regenerating *hydra*. Stabilized cytosolic actin structures disappear during the *de-novo* polarization, while polymerized microtubules remain. In our observations applied drugs that depolymerize actin filaments accelerate the symmetry breaking process, while drug-stabilized actin filaments prevent it. Nocodazole-depolymerized microtubules prevent symmetry breaking, but regeneration can be rescued by the microtubule-stabilizing drug paclitaxel at concentrations where microtubular structures start to reappear. We discuss the possibility that mechanical fluctuations induce the orientation and position of microtubules, which contribute to *β*-catenin nuclear translocation, to increase the organizer-forming-potential of the cells. Our data suggest that in regenerating *hydra* spheroids, microtubules play a pivotal role in the cooperative polarization process of the self-organizing *hydra* spheroid.

## 1 Introduction

The freshwater polyp *hydra* consists of a cell bilayer with a radially symmetric body, with one mouth opening. Hydra has astonishing capabilities for regrowth of lost body parts. Dissociation into single cells, mixing and random reaggration gives rise to a new hydra that forms from the disorganized heap. The first step towards regrowth from such a random cellular aggregate is the formation of an isotropic, hollow spheroid of about 10, 000 *hydra* cells. These then polarize and transform into fully functional *hydra* ^1^, albeit smaller than the adult consisting of about 100.000 cells. Hydra represents a well-known model organism for its remarkable regenerative properties ^2^. Regrowth from a spheroid requires *de-novo* axis definition ^3,4^.

Since the initial suggestions by Turing ^5^, Gierer and Meinhardt ^6^ discussed reaction diffusion mechanisms as a way to distribute morphogens that organize hydra regrowth and axis formation. To our knowledge molecular players could not be mapped to the reaction diffusion mechanism in spite of experimental efforts. The *dickkopf* gene has been suggested to play the role of an inhibitor ^7^ within the reaction diffusion framework. The evolutionary conserved Wnt3 *β* catenin Sp5 antagonist pathway ^8^ might be a candidate but further work would be required to confirm these ideas.

Fuetterer *et al*. described cyclic osmotic inflation and subsequent breakage of regenerating hydra spheroids. They showed an irreversible transition from symmetric to asymmetric inflation ^9^. The transition coincided with the emergence of at least one weak spot in the spheroid reducing the amplitude of the inflation. Soriano et al. ^10^ saw that with this reduction a weak temperature gradient could no more define the position of the future axis. Accordingly they proposed that this weak spot was a sign of early mouth formation, based on the timing and proposed molecular connections between WNT signaling and cell adhesion. Recently this idea was confirmed experimentally on biochemical grounds by Wang *et al*. ^11^.

Wnt and its associated pathways are involved in body patterning of cnidaria and bilateria. In *hydra* spheroids, *de-novo* head formation occurs via activation of the canonical Wnt pathway ^12–14^. The role of this pathway in the set-up of the head organizer and in the maintenance of the body plan of *hydra* has been investigated in great detail ^7,12,15–18^. The WNT expressing organizer is the first structure to be restored during *hydra* regeneration ^19^. In the absence of external stimuli, how and why the Wnt head-organizing centre is initially established at a certain location of the regenerating hydra spheroid is unknown. Strong electric fields can reversibly drive the evolution backwards from the adult to a spheroid, modulating Wnt3 activity ^20^.

*Hydra* axis establishment through WNT expression occurs at the moment where the gene expression pattern of the *ks1* gene, a marker of head-forming potential ^21^, scales in shape and size on the surface of the spheroid ^10^. Gamba *et al*. ^22^ theoretically showed that fluctuation-driven, nearest-neighbor based synchronization can lead to avalanche-like propagation of gene expression patterns that reproduce the experimental findings in space and time quantitatively. This is remarkable since the only free parameter of the mathematical model is the probability of information loss (in the percentile range) when synchronization between neighboring cells occurs. The theory only requires a hysteresis between cell states, independent of further details of intercellular communication. Nearest-neighbor communication as a means to generate asymmetry agrees well with previous suggestions by others ^23–25^.

The creation of asymmetry from the mechanism above requires that an asymmetric fluctuation be locked. This can be the result of an avalanche that comprises a substantial number of the cells. Given this assumption the model predicts that in a weak temperature gradient the position of the largest avalanches will depend not only on the direction of the gradient, but also on its strength or amplitude. This is in excellent agreement with experimental observations ^22^ and seems difficult to explain otherwise.

The mathematical model by Gamba *et al*. is based on a framework leading to a critical state ^26^. A critical state is characterized by the property that any perturbation is felt globally across the spheroid. This is due to tight coupling among neighbors that leads to the absence of a characteristic scale on which the information could be lost. As a consequence a critical system will respond to the smallest perturbation. In the case of hydra this means that the spheroid will pick up almost any kind of small initial asymmetry. This asymmetry will statistically propagate during further development. Observations that actin ^27^, WNT signaling ^28^, a temperature gradient ^10^, or maybe even gravity ^29^ can be responsible for the position of head regeneration, taken together, strongly favor the idea of criticality. Conclusions from axis inheritance as drawn in ^27^ are not suitable to understand a mechanism that amplifies almost any kind of weak asymmetry.

The fact that during regeneration a hydra spheroid exhibits a clear transition from isotropic to bipolar distribution of cell elasticity ^9^ points towards a role of the cytoskeleton. Soriano *et al*. observed that the level of mechanical slowdown of the osmotic inflation of the regeneration corresponded to the observed delay in this transition, irrespective wether the slowdown had been caused by temperature, mechanical-, or biochemical means ^30^. This suggests that the cyclic inflation may well represent the driving clock of the symmetry-breaking process, again pointing towards an important role of the cytoskeletal elements. Soriano et al. ^30^ showed that a mechanically driven Turing model that may lead to asymmetry can fit all of the observations well. A similar idea was taken up by Mercker *et al*., who postulated a mechanobiochemical feedback loop between tissue stretching, resistance to stretching, and the behaviour of a head-defining morphogen ^31^. However, the exact role of the mechanical clock and its connection to symmetry breaking still remain hypothetical ^32 31^. Again Turing inspired models appear as plausible but difficult to map to current molecular knowledge.

Following the critical model by Gamba *et al*., the control variable or “clock” should gradually increase the tendency of the cells to spontaneously express genes towards organizer formation. Regarding hydra, this tendency depends on *β*-catenin transport into the nucleus ^33 24^. In excellent agreement with the mathematical model, *β*-catenin acts independently of position ^34^. It is upregulated before the emergence of WNT ^16,33,35^. Hydra only possesses a single type of *β*-catenin. It is present in the cell periphery and degraded in the cytosol ^35,36^. Intracellular transport to the nucleus is likely to depend on the cytoskeleton.

The *hydra* cytoskeleton contains large super-cellular contractile actin structures, known as myonemes ^37^. Actin fibers were suggested to maintain the polarity in *hydra* regeneration from tissue fragments that retain their original polarity ^27^. Microtubules have been studied in hydra in the past ^38,39^. Recent work ^40^ showed that microtubules are involved in length regulation of the adult. However, we are not aware that microtubules were addressed in the context of hydra axis establishment before.

At the same time microtubules play important roles in the establishment of polarity during embryogenesis of diverse model organisms. Microtubules are involved in the polarisation of individual cells upon varied stimuli ^41^. Brunet *et al*. showed mechanosensitive induction of nuclear translocation of *β*-catenin in zebrafish. This induction can be directly activated by mechanical stimuli, and it leads to mesodermal invagination without prior Wnt activation. With the inhibition of microtubule polymerization using nocodazole, or inhibition of myosin II using blebbistatin, *β*-catenin-dependent mesodermal invagination is suppressed ^42^. Moreover, microtubules are involved in the asymmetric transport of Wnt8 mRNA during induction of the embryonic axis in zebrafish ^43^. The importance of microtubules in the establishment of embryonic polarity has also been shown in *C. elegans* ^44^. Kwan *et al*. showed that in *Xenopus*, a sufficient mass of polymerized microtubules is necessary for convergent extension during embryonic development ^45^. Non-canonical Wnt signaling and *β*-catenin signaling are involved in osteogenic differentiation upon oscillatory mechanical stimulus ^46^.

At a more general level, mechanotransduction is a means to propagate information at a large scale. An example is tissue elongation in *Drosophila* follicle cells that requires oscillating contractions of a basal actomyosin network ^47^. Mechanotransduction transmitted by the cytoskeleton can have a major role in gastrulation ^48,49^. The embryonic elongation of *Caenorhabditis elegans* involves myosin II ^50^ contraction. Mechanical fluctuations at a subcellular level can affect the polarisation of individual cells during development ^51–53^.

Here we investigate the role of the polar cytoskeletal filaments tubulin and actin in shape-symmetry breaking of regenerating *hydra* spheroids. The situation in hydra is different from other studied model systems since all the cells of a regenerating hydra spheroid can potentially express Wnt to become part of the organizer. At the same time, due to inflation, all the cells are submitted to the same mechanical strain. How irreversible asymmetry emerges from this isotropic situation is not known. We find that polymerized microtubules appear as a requirement for symmetry breaking with filamentous actin a putative antagonist.

## 2 Results

### Shape changes in regenerating *hydra* spheroids identification of symmetry breaking

Fragments cut out of the *hydra* body column fold inwards until hollow spheroids emerge, made up of a cell bilayer. Sufficiently small fragments loose their former body axis, if properly prepared ^10^. For the present study, these hollow spheroids were kept in drops of buffer that were hanging from the lids of petri dishes. The shapes of the regenerating small fragments were recorded as a function of time using a microscope equipped with a digital camera.

During the initial oscillations of the spheroids, cells were disgorged at deflations, which resulted in reductions in the diameter. (Supplementary Information, Fig. S1)). From the images obtained, the volumes of the spheroids were approximated by fitting an ellipse to the mid-plane sectional image and computing the volumes of the rotational ellipsoids (Fig. 1a). The regeneration patterns initially showed large oscillations in spheroid volume that decreased in amplitude at one point ^9^, to eventually stabilize. Very few markedly different patterns occurred (Supplementary Material Fig. S2), however, we did not understand how to quantify such deviations. Moreover, we believe that these were due to lacking adhesion or remaining asymmetry from the preparation. These (highly unusual) cases were discarded from further analysis.

**Fig. 1.**
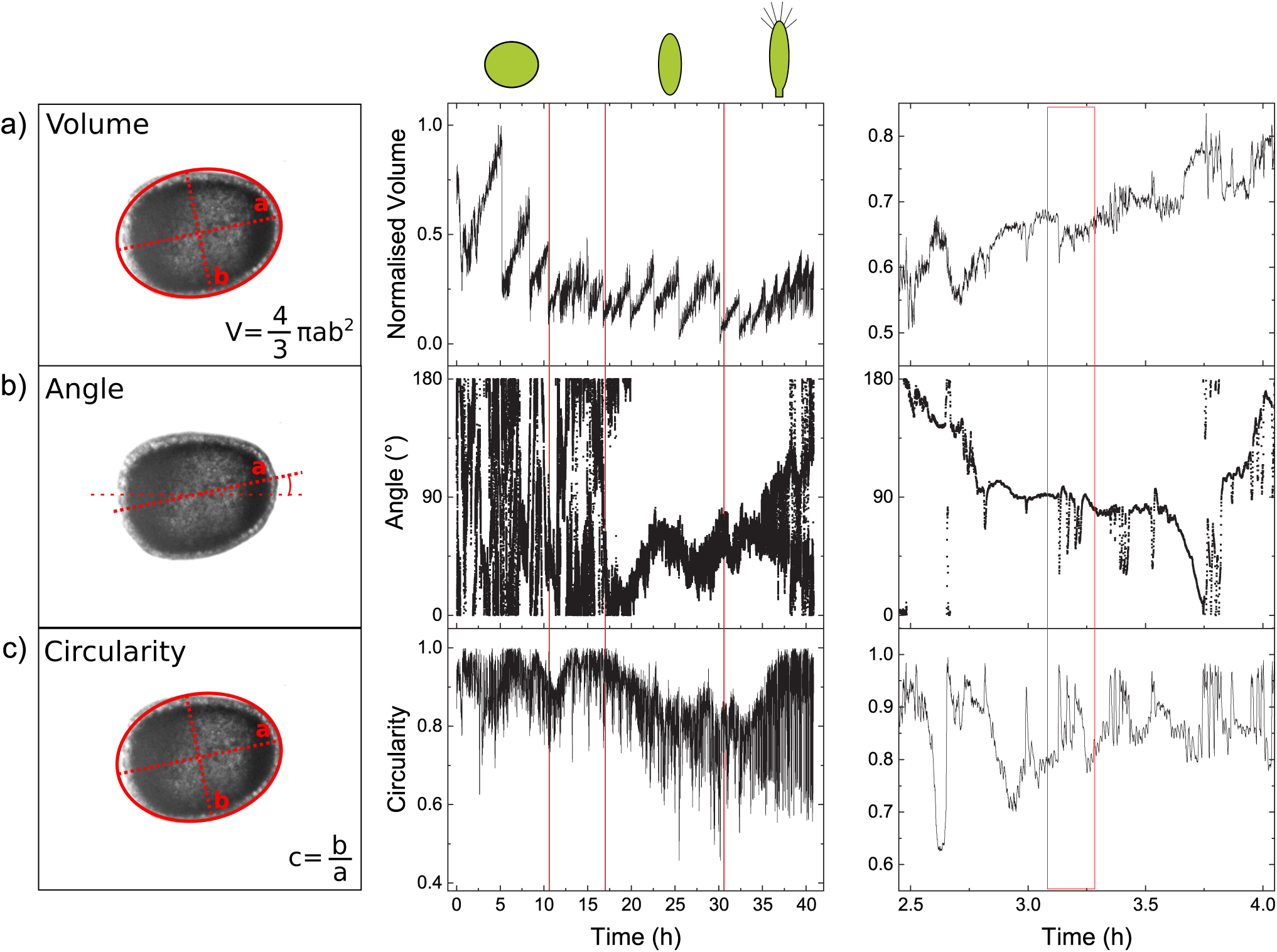
*Hydra* regeneration. Definition of the parameters (left) and their evolution as a function of time (centre) for a typical *hydra* regeneration (∼ 2 days). The pictograms (top) illustrate the overall shapes of the regenerating *hydra* during different periods of regeneration. **Left:** Images of regenerating *hydra* cell spheres were fitted to an ellipse. **(a) Volume**, as determined from the ellipsoid of rotation. **(b) Angle**, between the major axis of the ellipsoid and the reference axis (image frame). **(c) Circularity**, the ratio of the minor to the major axis of the ellipsoid, where a value of 1.0 corresponds to a sphere, and 0 to a straight line. **Centre:** The three parameters, Volume (normalized, arbitrary units), Angle (degrees), and Circularity as functions of time. The first red line indicates the end of the size-reduction period. In all studied cases shape symmetry breaking was only observed beyond this point. The second line indicates the transition from a situation where the sphere fluctuates by elongating in different random directions, to a situation where the axis remains irreversibly fixed for the sphere. This point defines the shape-symmetry breaking instant. The slow change in the angle after this point is due to slow rotation of the elongated cell sphere in the hanging drop. The circularity starts to more or less monotonically decrease almost at the same instant as the angle stabilizes. The ellipsoidal deformation becomes more and more pronounced in a given direction. The third red line indicates the moment where the *hydra* tentacles appear. After this point, the *hydra* moves actively and an ellipsoid does not fit the observed shape any more. At this point the values of volume, angle and circularity start to increase or scatter, but this does not represent quantitative changes. **Right:** Oscillations in the volume, angle, and circularity at a time scale of minutes to hours. An example of correlations between the volume, angle and circularity oscillations is highlighted by the red frame.

The angle of the major axis with respect to the image frame enabled us to follow the time course of the spatial direction of the hydra ellipsoids (Fig. 1b). Before symmetry breaking, on a timescale of minutes, phases where the angle remained stable in space alternated with phases where the angle varied rapidly (Fig. 1b, right panel). This variation was due to successive elongation of the spheroid in different spatial directions. The slow change in the angle, after the symmetry breaking, seen in figure 1b corresponds to a slow rotation of the spheroid in the hanging drop.

We took the circularity as the ratio of the minor to major axes of the ellipsoid, as described in figure 1c. A ratio of 1.0 corresponds to a sphere, and 0 corresponds to a straight line.

Irreversible axis establishment (*i*.*e*. broken symmetry) was identified by i) volume oscillations with diminished amplitude compared to the initial inflations (as in ^9,10^), ii) the irreversible transition to a stable direction of the axis in space and iii) the onset of a persistent decrease in circularity.

In figure 1 (right panel) fluctuations of volume, angle and circularity appear correlated. The cross-correlations among these parameters as a function of time were determined after subtraction of the underlying linear increases in the ‘sawtooth’ volume oscillations (Supplementary Information figure S4 for details on the subtraction). The rhythmic oscillations in the volume from the start until axis establishment, on a time scale of minutes, correlated with the oscillations of the orientation of the major axis (correlation coefficient, volume and angle: 0.9 ± 0.0306, *n* = 17 *hydra* spheroids). The time evolution of the angle, from the beginning of the first inflation until appearance of the tentacles, correlated with the circularity (correlation coefficient, angle and minor/major axis ratio: 0.83 ± 0.19, *n* = 17 *hydra* spheroids).

### Application of a mechanical stimulus using a micropipette

To apply a mechanical stimulus during the symmetry-breaking process, the spheroids were held in place using a micropipette (see Fig. 2, inset). There were no apparent changes regarding the regeneration time. The internal suction pressure of the pipette was maintained constant during the regeneration process. The stimulus from the pipette led to the heads emerging within an annular zone around the pipette tip at an average angle smaller than perpendicular to the axis of the pipette (Fig. 2, supplementary material Fig. S3). Three heads were pointing straight upwards, no heads were pointing downwards.

**Fig. 2.**
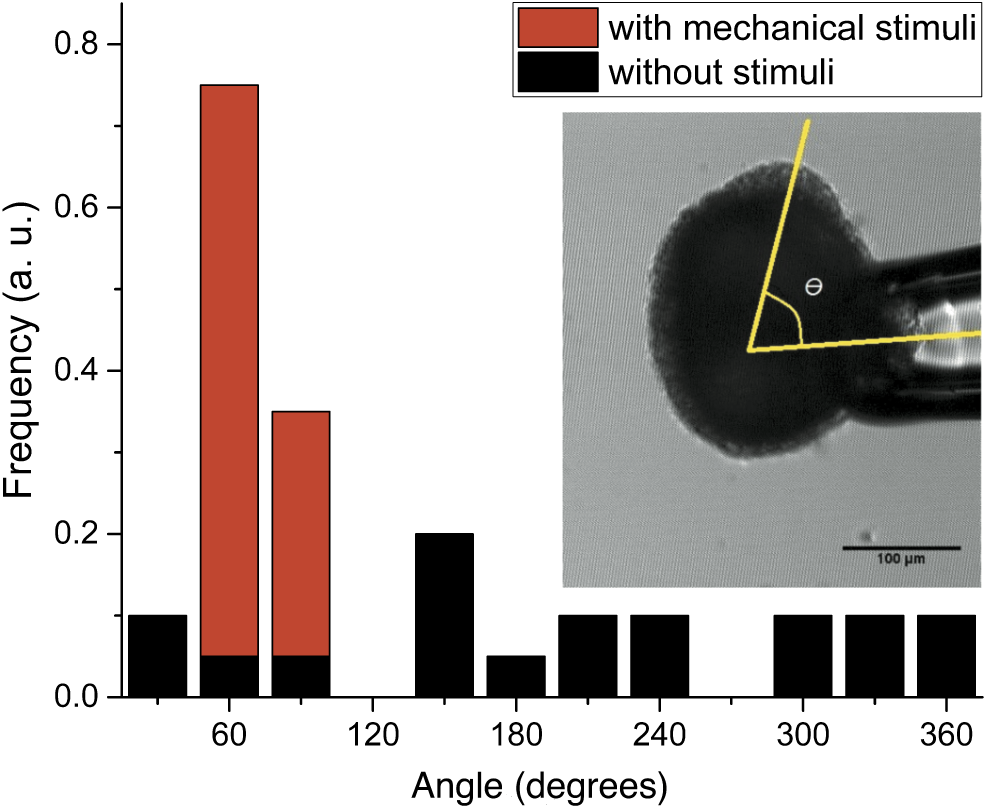
The mechanical stimulus of micropipette suction determines axis orientation. The distribution of the angles of *hydra* axis formation with respect to the pipette axis (‘with mechanical stimuli’) and in the absence of aspiration (‘without stimuli’), at the same spot but on the bottom of the culture dish. **Inset:** A *hydra* spheroid fixed by micropipette aspiration. The tissue was aspirated into the pipette with 60*µ*m in diameter. We observed that the *hydra* axis developed its tentacles in an area oriented at 40 − 70°to the pipette as indicated by the yellow lines.

### Actin structure under confocal microscopy

In adult *hydra*, the actin myonemes are arranged into two orthogonal layers. The inner layer surrounds the *hydra* in directions perpendicular to the body axis, and the outer layer spans the *hydra* parallel to the direction of the body axis (Fig. 3). After rounding up, in the early spheroid cortical actin and cytosolic actin stress fibers can be observed (Fig. 4). Both the actin myonemes and stress fibers tend to disappear as can be seen already within 60min after spheroid formation (figure 4). The actin myonemes and cytosolic structures are reestablished only after the shape-symmetry breaking, 18h to 24h after spheroid formation. This does not apply to spheroids that maintain their axis and do not lose the organization of actin.

**Fig. 3.**
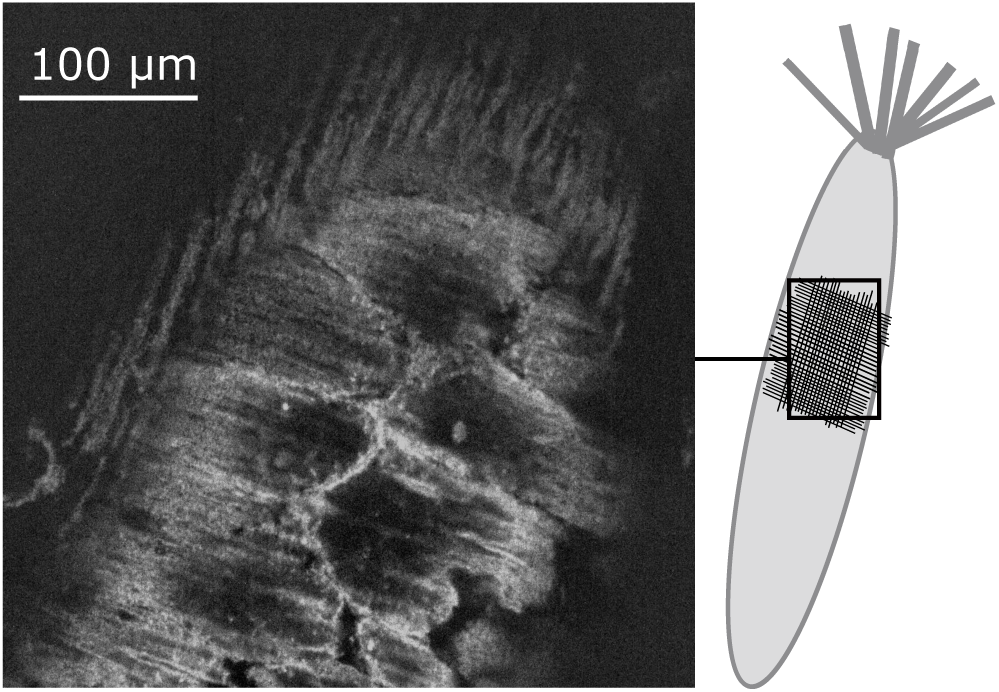
Actin myonemes in adult *hydra* as observed by confocal microscopy. The cortical actin and the actin myonemes were stained using rhodamin-phalloidin. The orthogonal layers of the actin myonemes can be distinguished. In regenerating, axis forming spheroids we could not observe the equivalent structure.

**Fig. 4.**
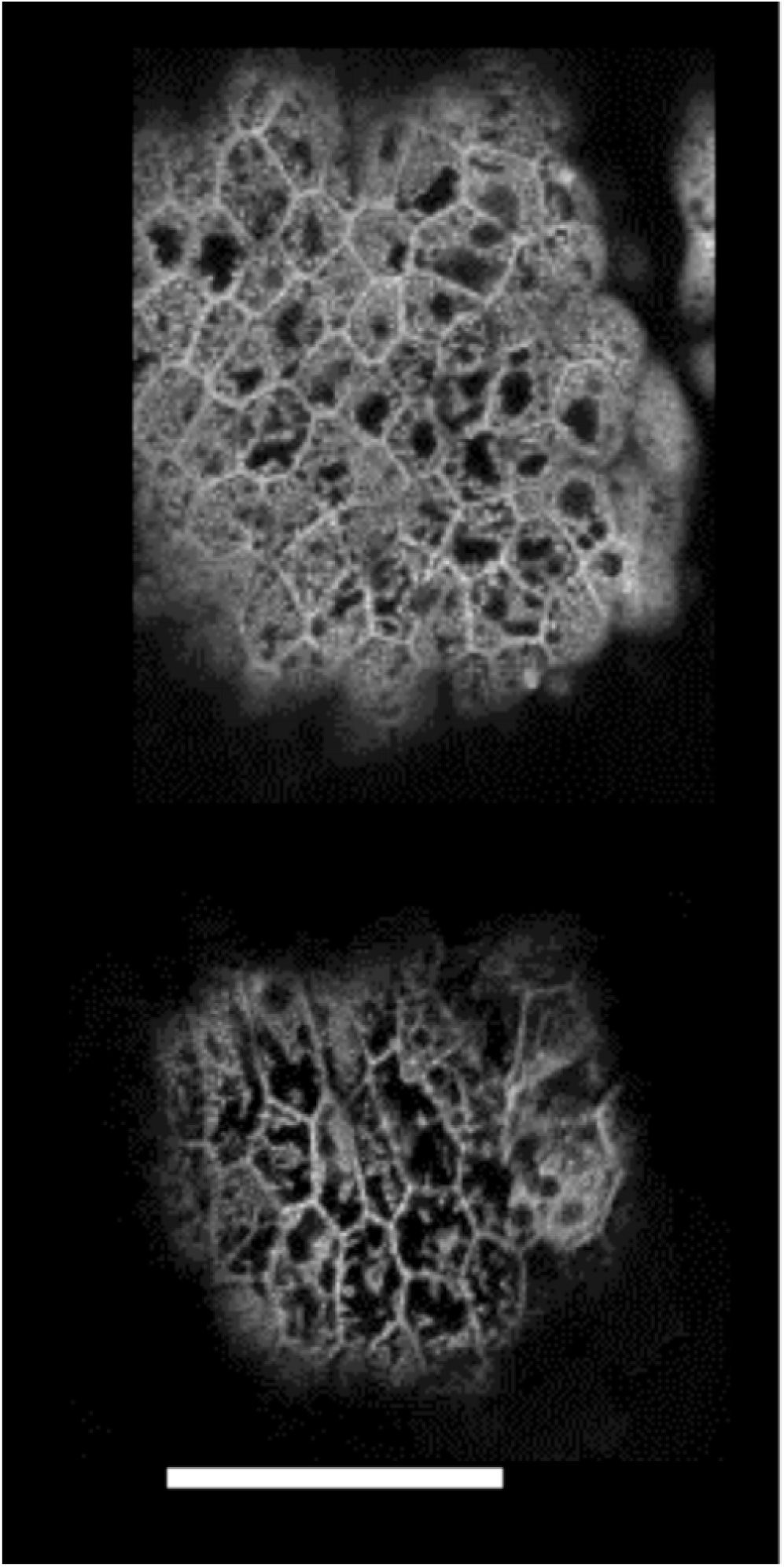
Loss of intracellular actin structures in transgenic live actin-GFP *hydra* spheroids observed by confocal microscopy. Representative images of the ectodermal layer taken 60min after a fragment was cut from a hydra for regeneration. Scale bar is 50*µ*m **Top:** *Hydra* spheroids with an initial diameter of ≈ 500*µ*m. The cellular actin structures are visible as a network. This spheroid showed asymmetric fluctuations in a fixed direction from the beginning. **Bottom:** Smaller spheroid with an initial diameter of ≈200*µ*m. The intracellular structures start to disappear, but within the cellular cortex actin structures appear unchanged. This type of spheroids underwent symmetry breaking as described in figure 1.

### Influence of cytoskeleton-directed drugs on *hydra* spheroid motion

We investigated actin structure-modifying drugs for their impact on axis formation in *hydra* spheroids. Phalloidin (actin stabilizer), Y-27632 (inhibitor of p160 ROCK and actin contractility), and SMIFH2 (inhibitor of Formin and *de-novo* stress fibre formation ^54,55^), as well as the microtubule-modifying drugs nocodazole (depolymerizing) and paclitaxel (stabilizing) were used. The influence of the drugs on the evolution of the spheroid shape is shown in figures 5 (actin-modifying drugs) and 6 (microtubulemodifying drugs). Using brightfield microscopy we could not identify by eye any systematic changes in the individual cell shape or volume that were linked to the application of drugs. Nocodazole-treated spheroids did not develop at concentrations as low as 0.1*n*M (*n* = 15). However, the development was rescued by addition of 0.1*µ*M paclitaxel 24h later (14 out of 15 spheroids) (Fig. 7). Microtubule staining revealed that the rescued development correlated with increased presence of polymerized microtubules including super-cellular structures, re-emerging after 24h, that is, before axis formation (Fig. 8).

**Fig. 5.**
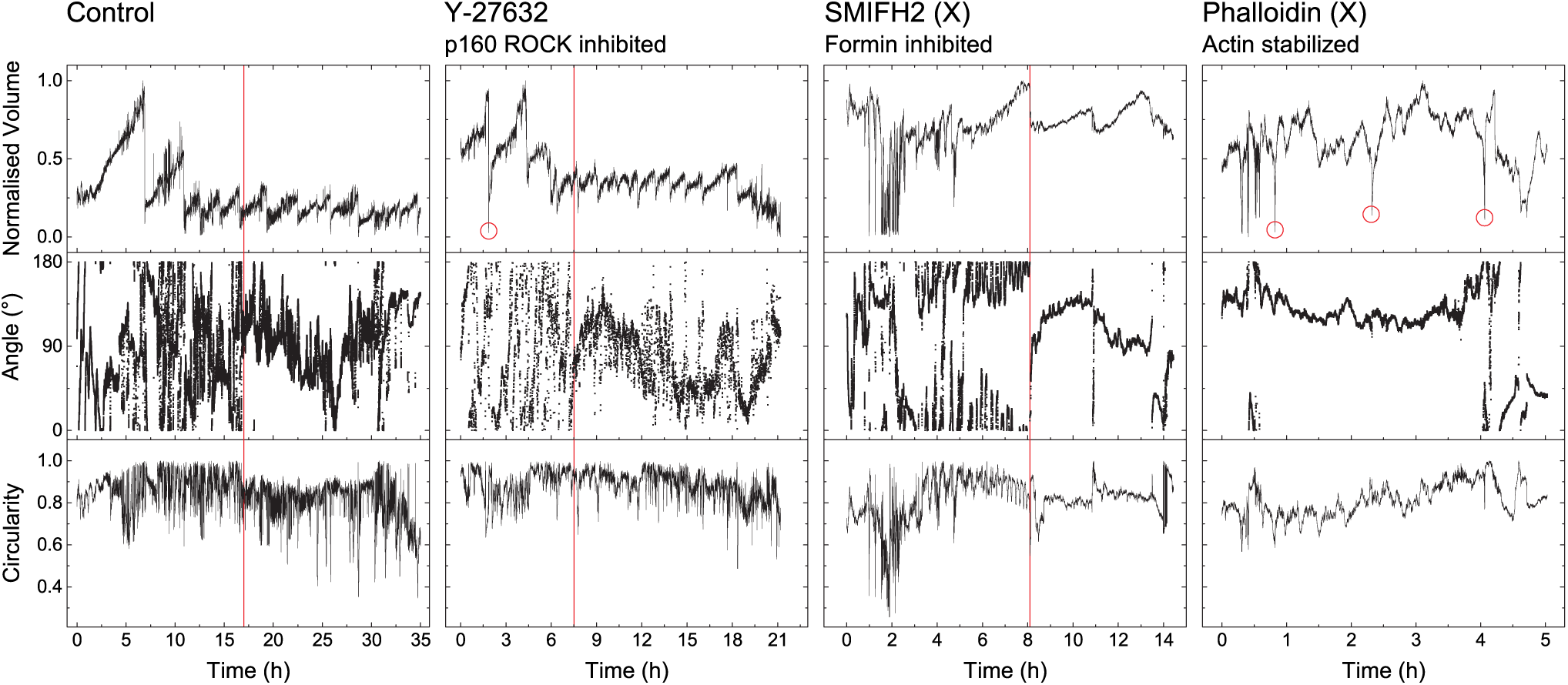
Normalized volume, angle and circularity of regenerating *hydra* spheres under treatments with actin-modifying drugs as a function of time. Not completed regenerations (in the cases of SMIFH2 and Phalloidin treatment) are indicated by (X). **Control:** Untreated spheroid. **Y-27632** (ROCK inhibitor): Inhibition of actin contractility. The initial volume inflations showed long, downward-pointing peaks that were not present in the control (red circles). The symmetry breaking appeared much earlier (red line), ∼7.5h versus ∼17h in control. However, the axis angle appeared more scattered compared to control, indicating more pronounced fluctuations of the shape. **SMIFH2** (Formin inhibitor): Inhibition of *de-novo* stress fibre formation. The sudden stabilization of the spatial orientation of the major axis remained (red line), indicating formation of a persisting axis. Beyond this point the spheroids failed to elongate further along the established axis and did not regenerate. **Phalloidin** (actin filament stabilizer): The large volume sawtooth oscillations seen in the control did not occur. Downward pointing peaks of the volume appeared (indicated by circles). The axis angle remained stable in space during the entire observation time, which indicated that the initial symmetry was not lost. At the same time, the cell balls rounded up, as can be seen from the increase in circularity. Eventually the spheres started to disintegrate.

**Fig. 6.**
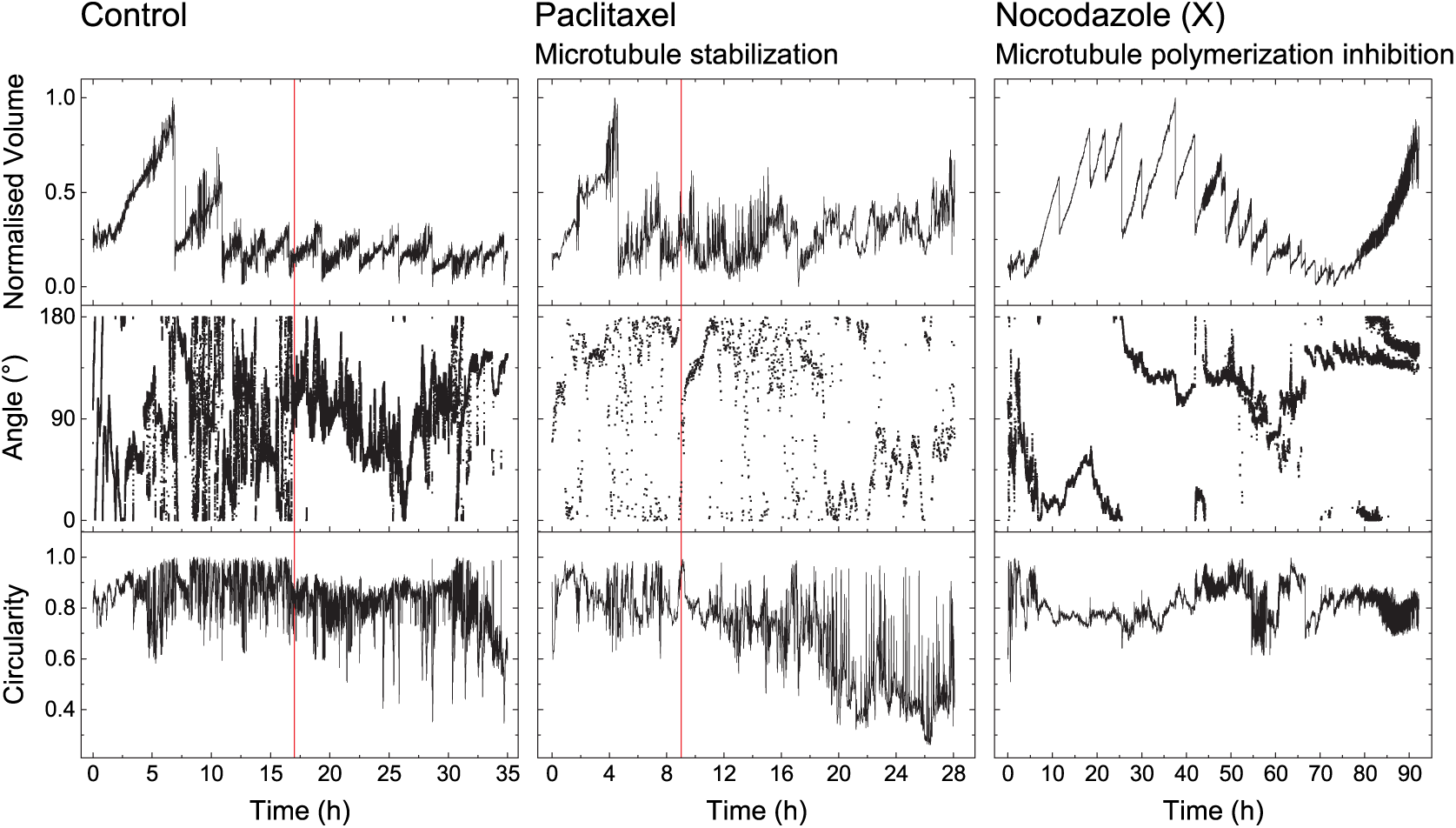
Normalized volume, angle and circularity of regenerating hydra spheroids as a function of time and treatment with microtubule-modifying drugs. Unsuccessful regeneration (in the case of Nocodazole treatment) is indicated by an (X). **Control:** Untreated spheroid. **Paclitaxel** (microtubule filament stabilizer): The red line indicates the moment where the sphere started to elongate irreversibly, and the direction of the axis was established. Although the direction of the axis showed more pronounced fluctuations in time than the control, the *hydra* regenerated successfully. **Nocodazole** (microtubule polymerization inhibitor): The axis direction and circularity during the first 24h suggest that the symmetry of the spheroid appeared broken from the beginning. After 24h, the spheroid started to shrink and then disintegrated as reflected by the increase in volume starting at ∼ 75h.

**Fig. 7.**
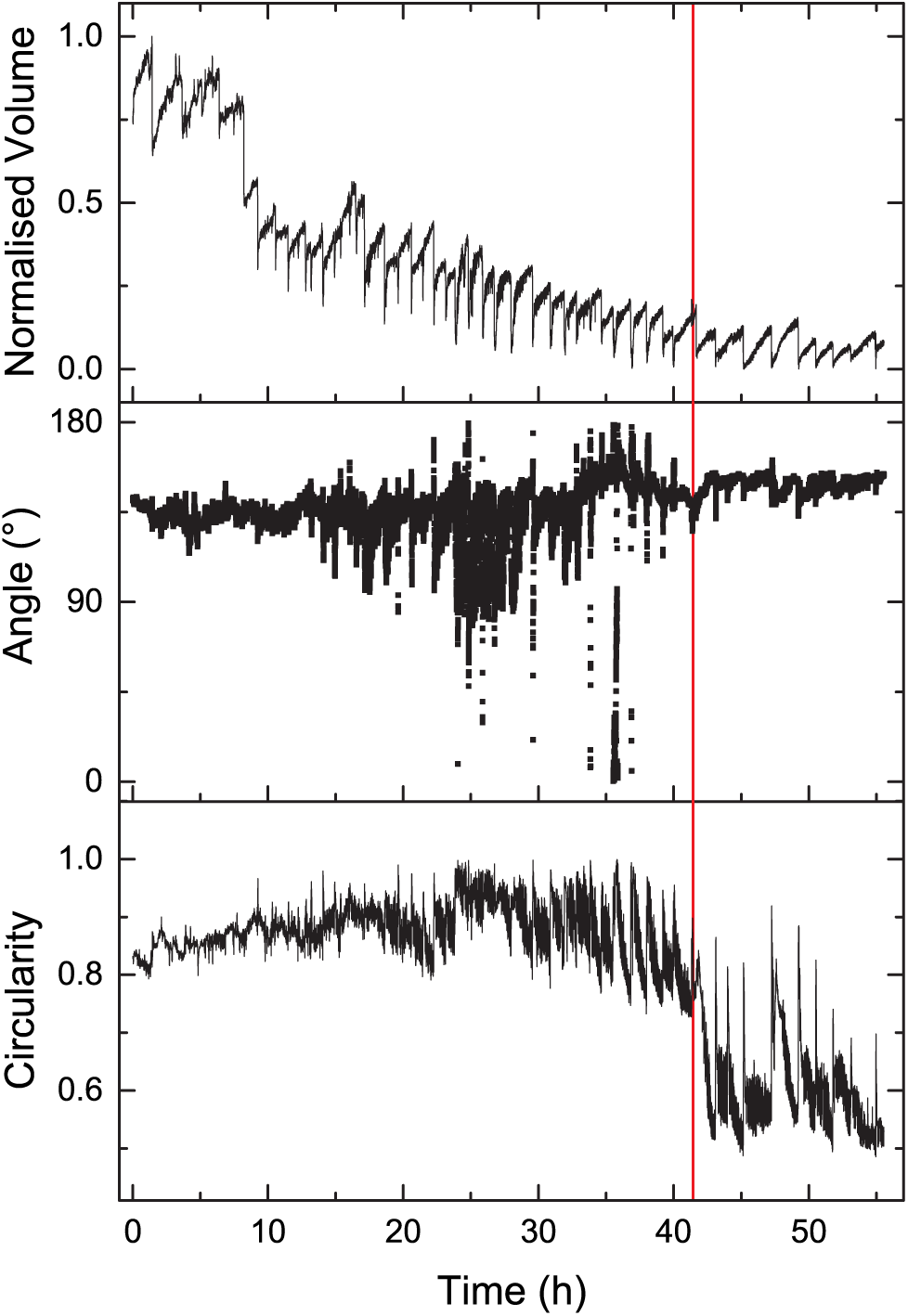
Shape changes of a *hydra* spheroid treated for 24h with 0.1nM nocodazole, then exposed to 0.1*µ*M paclitaxel. The *hydra* spheroid established an axis only after about ∼41h (red line) counted from paclitaxel addition, that is, 65h after spheroid formation. Nocodazole-treated spheroids without paclitaxel treatment did not show signs of developing asymmetry and disintegrated (Supplementary Information, Fig. S6)).

**Fig. 8.**
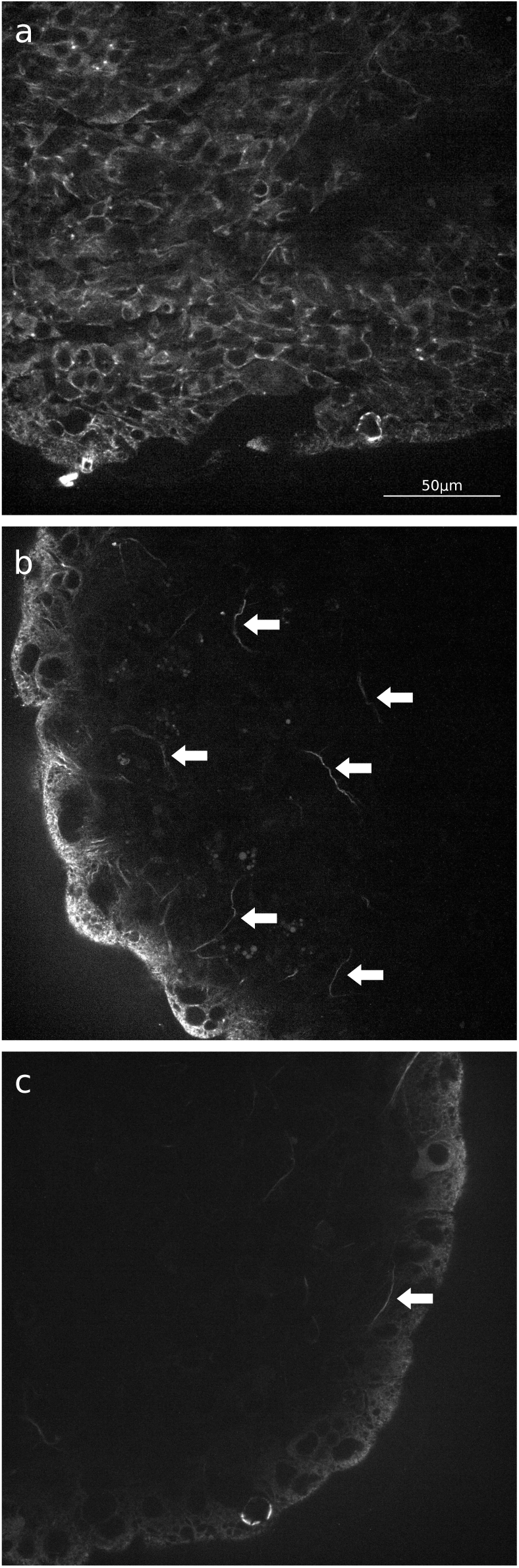
Spheroids stained for microtubules. The spheroids were fixed, stained for microtubules and observed under the confocal microscope. **(a) Control**, stained 24h after spheroid formation. **(b) Paclitaxel** rescued spheroids, stained 24h after paclitaxel treatment: there are polymerized microtubules, which appeared as supra-cellular structures (arrows) that were about 50*µ*m long; however, there was less polymerized tubulin than in the control (a). **(c) Nocodazole**-treated *hydra* spheroids, stained simultaneously with (b): no polymerized microtubules were seen, except for one supra-cellular structure (arrow). The images are at the same scale; scale bar in (a)

In figure 1 (right panel,b) periods of quiescence can be distinguished from periods where fluctuations of the angle occur. We chose a minimum threshold amplitude to define fluctuations, and plotted the summed probability of periods of quiescence as a function of the duration of the periods of quiescence (see Image analysis in the methods section). A log-log plot reveals that paclitaxeltreated (stabilized microtubules) regeneration had a similar tendency to a power law as the control; however, this occurred with reduced absolute probabilities (Fig. 9). This indicates that paclitaxel increases the likelihood of shape fluctuations, in good agreement with figure 6.

**Fig. 9.**
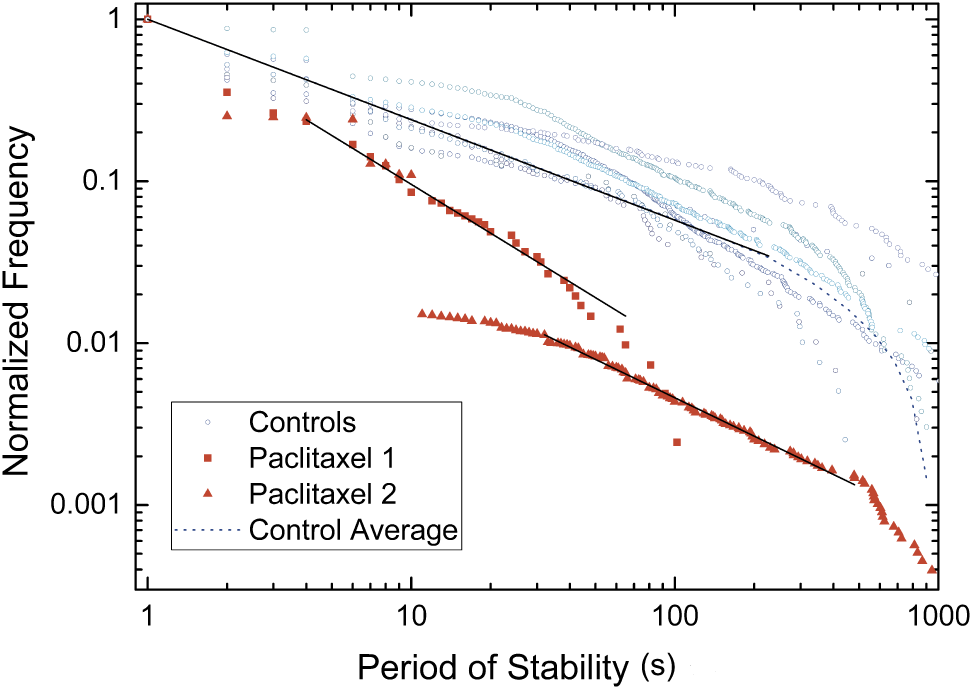
Frequency (or probability) of occurrence of a fluctuation above threshold as a function of the inter-fluctuation time interval. The control was approximated by a power law of *y* = *c · x*^−*β*^ with *β* = 0.62 ± 0.003. Treatment with the microtubule-stabilizing drug paclitaxel shifted the distribution to lower values. The exponents remained comparable: *β* = 1.00 ± 0.02 and 0.79 ± 0.01, for the two paclitaxel cases.

Fourier transform of the volume (Fig. 1 a, centre) revealed 1*/ f* scaling behaviour of the frequency spectrum, 1*/ f* ^*α*^, where *α* = 1.43 ± 0.21 (*n* = 8) for control, above frequencies of 0.01Hz (Supplementary Information, Fig. S4 and S5). Upon addition of nocodazole the amplitude was reduced by a factor of 10. The mean exponent in *hydra* treated with nocodazole was 0.84 ± 0.12 (*n* = 9 regenerations) corresponding to a broader distribution in frequencies.

## 3 Discussion

*Hydra* tissue fragments tend to fold in to eventually form hollow spheroids made of a cell bilayer. If properly prepared, tissue fragments beyond a certain size undergo de-novo axis definition. That case clearly differs from hereditary axis formation ^10^. For small spheroids that undergo *de-novo* axis formation, we identified transitions from a state of elongation in arbitrary directions to a state where the fluctuations followed a transient axis. We consider that this case reflected symmetry, since the situation was reversible. This corresponds to the definition applied to the distinction between symmetry and asymmetry in physics, which has proven sensible in most complex problems. The abruptness of the last, irreversible transition to an increasingly elongated state not only correlated with the irreversible axis positioning in space, but also with the changes in volume inflation that appeared shortly beforehand. This behaviour exhibits mechanical synchronization of the entire spheroid at the symmetry breaking instant, suggesting supra-cellular cytoskeletal organization. It is, to our knowledge, the first spheroid spanning manifestation of axis formation that occurs during regeneration. At this stage, a temperature gradient is no longer able to direct the emerging axis ^10^. At the same time the irreversible change in all three parameters, volume, circularity, and axis direction makes irreversible axis formation simple to detect.

There are other criteria to decide about successful axis establishment. In many studies the axis is taken from the position of emerging tentacles, This is not applicable here, since drugs may well preclude tentacle formation without preventing, however, the observed symmetry breaking and axis formation that occurs substantially earlier. Another possibility would be to consider the spot where WNT expression occurs as a first signature of symmetry breaking. However, it may be that early organizer formation is reversible ^20^, in particular early molecular asymmetries may not always translate to axis establishment. Moreover, for our study we only need to know if an axis is irreversibly established at some point or not. As a consequence, for our purpose molecular asymmetries are neither more nor less suited to study symmetry breaking than any other manifestation of asymmetry such as shape, in particular if the latter extends throughout the spheroid. Related considerations have been discussed earlier in a different context, leading to analog conclusions ^56,57^.

Freshly formed spheroids repeatedly inflate until they rupture, thereby losing cells. On theoretical grounds, osmotic inflation and tissue rupture have been suggested to be the driving forces for the resulting sawtooth oscillations in volume ^58^, in good agreement with the experimental findings. A dependence of the time span required for regeneration on the initial size of the spheroids was reported previously ^1,30,59^. In the present study, the timespan correlated with the time needed to decrease the size of a spheroid until a given size was reached (Fig. 1). The impact of osmosis scales as the surface/volume (*d*^2^*/d*^3^, where *d* is the diameter), which means that smaller spheroids inflate relatively faster, and this could be a way to sense the size of a spheroid. However, the time required for artificial stem-cell clusters to polarize without external cues ^60^ also depends on the tissue size, which indicates that there might well be other mechanisms involved here.

Within the inflation and rupture sawtooth oscillations, we identified fluctuations at a shorter time scale (Fig. 1, right) that hinted at active contraction and/or controlled relaxation against the osmotically inflating spheroid. Spectral analysis by Fourier transform followed 1*/ f* ^*α*^ power-law description (Supplementary Information figure S5). Note that a power-law description means the absence of a characteristic size, that is, scale free behavior, which entails unusual properties. The distribution of time intervals of the axis directional fluctuations (Fig. 9) also agrees well with a power law description. Soriano *et al*. observed *ks1* gene mRNA expression patterns that obeyed a power law description at the moment of symmetry breaking ^10^. As discussed in the introduction, a self-organized, critical ^61^, nearest-neighbor-coupled, lattice-based model quantitatively reproduced the scale-free distribution of *ks1* expression patterns as well as the axis orientation in a temperature gradient. The model was robust to the choice of its sole free parameter, the information exchange probability between neighboring, critically excited cells, fixed to about 0.95^22^. Critical behaviour is observed in many complex, dissipating systems ^62^, in particular in the biological context ^63^.

### Actin depolymerizing drugs speed up the definition of axis position

We treated the regenerating spheroids with with the ROCK inhibitor Y-27632. Fibroblasts treated with Y-27632 showed reduced myosin light chain phosphorylation and increased twodimensional spreading areas, with reduced levels of stress fibers at the cell centre. This was accompanied by increased actin concentration at the periphery; however, the line tension was decreased simultaneously ^64^. Supposing the same action in *hydra* cells, the effects of the drug relate very well to the observed changes in inflation patterns. In general, decreased actin stress fibre levels reduce integrin-mediated cell adhesion. This decreased cell adhesion agrees well with the increased tendency to rupture, which leads to the observed diminished amplitude (and increased frequency) of the sawtooth inflation-relaxation patterns. In conjunction with the reduced contractility of the cell periphery, the increased surface area of Y-27632-treated spheroids matched well the overshoots during relaxation (Fig. 5, red circle) of the spheroid volume: As the spheroid breaks up due to osmotic inflation, there is a strong overshoot to low volumes that quickly disappears, which is again a sign of increased cell surface area.

We treated the regenerating hydra spheroids with the Formin homology 2 domain inhibitor SMIFH2. Its precise mode of action remains poorly understood ^65^. Nevertheless in *in-vitro* studies, SMIFH2 diminished f-actin levels during the first hour(s) of application in different cell types. Subsequently, the entire cytoskeleton is reorganized, although with a notably different structure. Our observations in regenerating *hydra* reflect this type of behaviour: initially there was no osmotic inflation, a likely consequence of the loss of f-actin and the resulting weak cell adhesion. Oscillations were rescued after about 4h, which we interpret by the reappearance of f-actin. The volume then reaches a maximum compared to the control in terms of amplitude and time of occurrence.

We note that for both actin depolymerizing drugs the observed inflation pattern are in excellent agreement with our expectations from the literature. From the observed pattern we only conclude that the drugs seem to work in similar manner in hydra cells as in the other cell types that they were studied in.

It appears that although the pattern of motion is substantially modified by the application of actin depolymerizing drugs, this does not prevent the symmetry breaking process. We note that the asymmetry occurs much earlier than in the control (8 vs 18 hours from start of inflation) irrespective of the nature of the actin depolymerizing drug. In ^30^ an extensive study on nearly 100 regenerating spheroids showed that the shortest time-span until early mouth formation varied by only 10 %. Accordingly our observation is significant. Due to the different working principles of the drugs this points towards actin loss as the main reason for this change.

Phalloidin stabilizes actin fibers, to prevent the reorganization of the actin cytoskeleton and stiffen the cells. Accordingly the sawtooth oscillations observed here were of low amplitude (Fig. 5). A high oscillation frequency with low amplitude from the beginning of the inflation process indicates either frequent pressure release or residual contractility. Phalloidin-treated spheroids rounded up, which indicated the loss of any residual asymmetry in the shape, until they eventually disintegrated after 2 days. We did not detect any symmetry breaking in the presence of phalloidin.

We observed that intracellular actin structures (Figs. 3, 4) weakened before symmetry breaking. Cortical actin, however, remained present even during the symmetry breaking process, and may still contribute to a mechanotransduction process. In our experiments, larger fragments formed spheroids that remained polarized throughout the regeneration process and maintained their initial actin organization (Fig. 4). This is in good agreement with ^27^, who showed that the super-cellular orientation of actin fibers provided the direction for the body axis in spheroids that retain their former axis. We suggest that the presence of actin filaments slows or prevents de-novo axis formation.

### Microtubule polymerization modifying drugs reveal polymerized microtubules a prerequisite to axis formation

In traction-force rheological experiments on fibroblasts that were non-specifically glued to opposing surfaces, neither the microtubule-modifying drug nocodazole nor taxol had any noticeable influences on the mechanical responses of the cells to deformation ^66^. In contrast, an actin-modifying drug resulted in large changes. Although microtubules do not directly contribute to the production of cytoskeletal forces, they are well known to orchestrate cell mechanics in a biological setting. In particular, the depolymerization of microtubules stimulates cell contraction via Rho and subsequent activation of the actin cytoskeleton, which results in the stimulation of cell adhesion (see ^67^ and references therein).

Here we observed that the nocodazole-treated *hydra* spheroids inflated more slowly compared to the control, and reached their maximum volumes only after as long as 36h (*i*.*e*., compared to 6h for the control). This behaviour is well explained by a more contractile actin cytoskeleton that yields more slowly against the osmotic inflation pressure than in the control. The volume of nocodazole-treated spheroids increased more on a relative scale (Fig. 6). This can be understood in terms of the highly contracted (and therefore smaller) spheroids used as the starting point. At the same time, the increased cell adhesion that comes with increased contraction ^68^ prevents rupture, so that the volume can be inflated more before rupture occurs. Nocodazole prevented *hydra* symmetry breaking and regeneration (Fig. 6).

We observed that the microtubule-stabilizing paclitaxel accelerated the rhythm of the high-frequency oscillations (Figs. 6 and 9). On the timescale of hours, paclitaxel has been shown to completely inhibit catecholamine release, thereby increasing cytoplasmic free Ca^++69,70^. In *Hydra attenuata* catecholamine inhibitors could prevent regeneration. Dopamine is a specific catecholamine where the inhibition induces morphogenetic anomalies in gastral spheroids, which do not retain their natural organizer ^71^. However, when paclitaxel was applied, no morphological anomalies were observed in any of the surviving spheroids in the present study, so this precise mechanism appears not to be important in our case. However, the activity of K^+^ channels is determined by cytoskeletal interactions ^72^. Even without superordinate neurological structures, the observed high frequency oscillations on top of the sawtooth oscillations may in principle result from interdependent effects of the cytoskeleton under osmotic tension in conjunction with cellular electrophysiology. We suggest that the same applies to the reduced fluctuations caused by nocodazole.

Paclitaxel increased the amplitudes of the sawtooth volume oscillations. An increased amplitude requires either increased adhesion within the tissue, or reduced cell contractility. The presence of microtubules is well known to reduce cell contraction controlled by Rho. In the presence of paclitaxel, the symmetry breaking event occurred earlier (around 10 h, instead of 18 h for control), which again is well explained by the lower levels of actin fibers that come with the stabilized microtubules.

Since drugs may act in different ways, we checked if nocodazole treated spheroids could be saved by exposing them to paclitaxel. Both drugs have different actions, however they overlap with respect to microtubules as their center of action. Indeed nocodazole above 0.1 *n*M was sufficient to prevent *hydra* regeneration (Fig. 6); however, application of microtubule stabilizing paclitaxel could rescue the process (Fig. 7). The amount of paclitaxel required to rescue nocodazole-treated spheroids correlated with the increased appearance of micro-tubular structures before symmetry breaking as observed in fluorescence microscopy (Fig. 8). We conclude that indeed microtubules are the functional elements that were modified by the drugs paclitaxel and nocodazole.

### Microtubules and actin are likely to act as antagonists with regards to axis formation

We see that actin filaments become weaker in control spheroids that undergo symmetry breaking (Fig. 4). Irrespective wether the actin filaments are removed through SMIFH2, or Y-27632, or by paclitaxel promoted microtubular growth, the emergence of asymmetry is accelerated by about the same time span. The fact that actin needs to be removed for *de-novo* axis formation is in excellent agreement with ^27^, who found that actin stabilizes asymmetry. The fact that the filament stabilizing drug phalloidin prevented symmetry breaking appears as a natural consequence. We suggest that actin filaments tend to maintain asymmetry, while growing microtubules tend to diminish actin filaments via Rho. At the same time, in the absence of actin, microtubules are the only macroscopic polar filaments, and they are required for axis formation. We understand that both types of filaments have antagonist action regarding *de novo* axis formation.

### A connection between Wnt expression, *β*-catenin and microtubular transport?

The current knowledge concerning the relevant Wnt molecular pathways and the cytoskeleton are illustrated in Fig. 11. The canonical Wnt pathways involve nuclear translocation of *β*-catenin, which in turn enables the affected group of cells to change their fate, and differentiate into a head organizer. During *hydra* axis-formation, increasing *β*-catenin nuclear translocation appears a natural way to slowly increase the head-forming potential as suggested in ^22^. In the absence of dishevelled, a dynein *β*-catenin complex can be expected to form in the cell cortex that can then actively move on microtubules in direction of the nucleus ^73^. If the microtubules extend from the cell nucleus to the cell periphery, the flux of *β*-catenin into the nucleus can be expected to increase. A sufficient increase here would enable a fraction of *β*-catenin to escape from degradation by the *β*-catenin degradation cascade in the cytosol so that the nucleus can be reached. The increase could be promoted by the observed cell stretching and relaxation oscillations, to make the mechanical oscillations appear as the clock driving the process ^30^. Another way to promote transport through the cytosol at increased rates could be the arrangement of microtubules to parallel bundles as a consequence of cell deformation (Fig. 10). Indeed, in adult *hydra* cells elongate in the direction of the microtubules, while they contract in the perpendicular plane ^40^. If the tissue is stretched by external forces, the microtubules will tend to orient in parallel to the stretch. This will promote further outgrowth and deformation towards a cylindrical shape during *hydra* spheroid inflation, thereby stabilizing elongated shapes. Indeed this type of deformation is observed (Fig. 10).

**Fig. 10.**
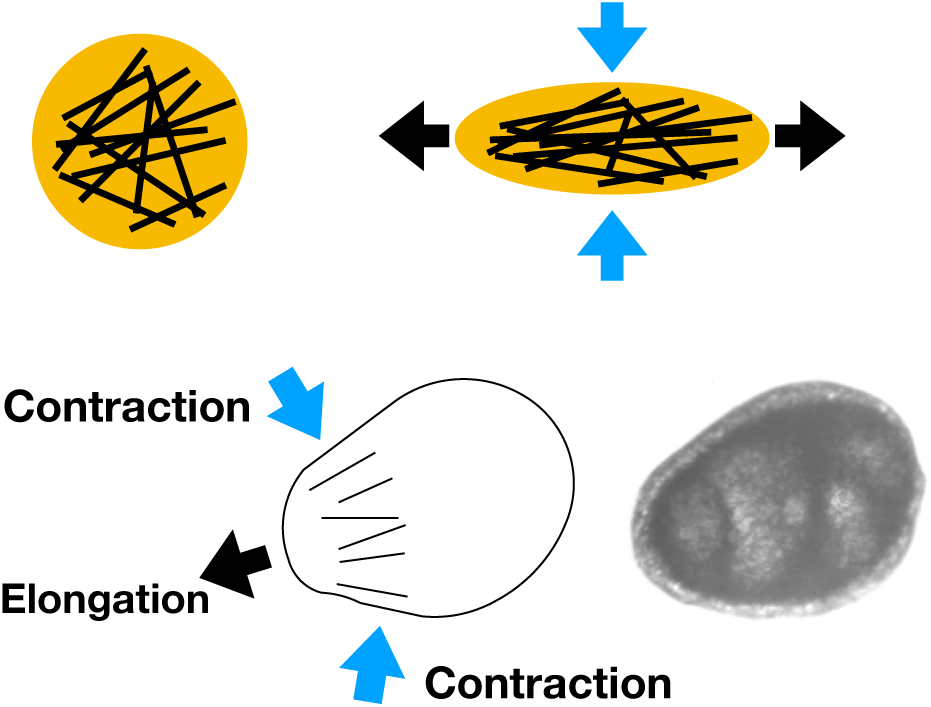
*Hydra* cell shape and microtubules. In adult *hydra*, microtubule alignment follows cell elongation in the microtubular direction ^40^. **Top:** As a cell (left) is stretched, the filaments (black straight lines) will tend to align in the direction of the stretch (right). **Bottom:** Following this mechanism, this can be expected to cause bulging (left) as observed during osmotic inflation of non-polarized hydra spheroids (right).

**Fig. 11.**
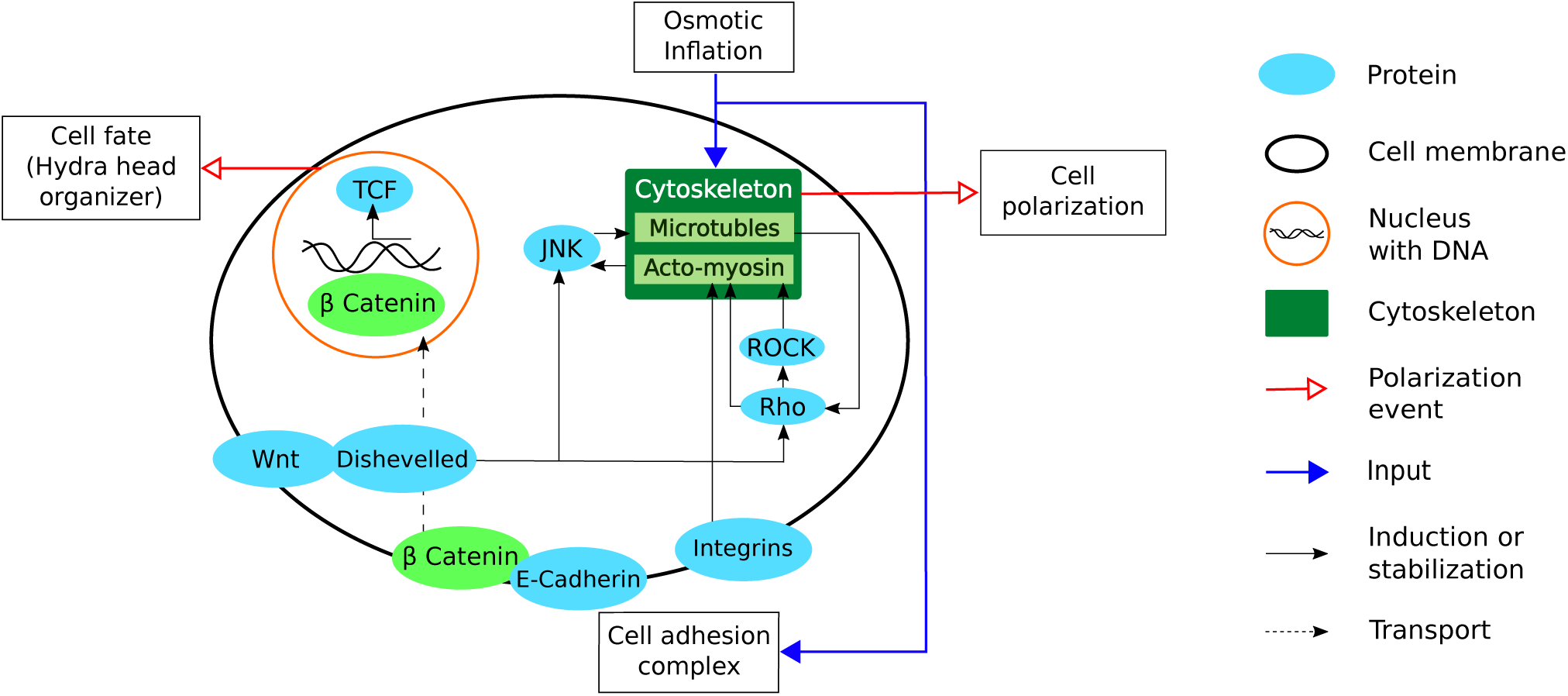
Wnt pathways overlap with the biochemical reactions to mechanical stimulus. The dishevelled complex protects cytosolic *β*-catenin from degradation by the catenin degradation complex. *β*-catenin, which is located in particular at the intracellular side of E-cadherin, can then be translocated into the nucleus to activate the transcription of T-cell factor (TCF) ^74^. TCF induces the formation of the head organizer in *hydra* ^13,15^. In *Drosophila* ^75^ and *Zebrafish* ^42^, *β*-catenin induces mesodermal invagination, which leads to gastrulation. In turn, *β*-catenin nuclear translocation can induce Wnt expression ^76^. Wnt controls acto-myosin contraction via Rho, including ROCK-dependent protein kinase ^77^. The Wnt-dishevelled complex/ pathway can act locally to stabilise microtubules ^78^. In addition, the Wnt-dishevelled complex can reorganize the microtubule cytoskeleton via JNK (c-Jun N terminal kinase, which is involved in cell polarization) ^79^. JNK-mediated cell polarization can also be induced via mechanical stimuli ^80^, thus leading to mechanically induced cell polarization. Wnt-controlled JNK activity mediates convergent extension during embryonic development ^81,82^. Within this framework, we hypothesize that in conjunction with long, oriented microtubules, a *β*-catenin - dynein - microtubule complex ^73^ can increase the transport of *β*-catenin to the nucleus.

This ideas above perfectly agree with experiments where a micropipette deformed the spheroid. The head indeed appeared within the annulus where the tissue is stretched most because of the suction so that cells are elongated in one direction in this area. Another interesting observation is that the heads tend to emerge in the upper part of the spheroid, clearly showing that either light or gravity determine the axis position in the non-perturbed direction in space. This again points out the critical nature of the symmetry breaking process that manifests in the observed scaling relations or power-laws as well as the pick-up of minor, external perturbations.

## 4 Conclusions

Symmetry breaking in *hydra* spheroids of around 10,000 cells exhibits coordinated mechanical fluctuations and oscillations. Here, we have shown that these spheroids show a clear transition to an oblong shape, from where on the position of the related axis remains irreversibly defined. Cytosolic actin fibers disappear before axis definition, and their removal by actin-modifying drugs accelerated the symmetry-breaking process. This agrees well with tissue fragments, where the remaining actin fibers dictate the polarity of the regenerating *hydra* ^27^. For de-novo axis formation these filaments need to be removed. We have presented strong evidence suggesting that polymerized microtubules are required for symmetry breaking. In excellent agreement with observations in other species, we suggest that microtubule orientation can increase *β*-catenin translocation in conjunction with dynein ^73^ in direction of the nucleus, which eventually results in the differentiation of a head organizer. The recent establishment of 3D *β*-catenin live microscopy ^83^ using fully transgenic *hydra* ^84^ could reveal very helpful in further studies. The mechanical connection through the extracellular matrix is a suitable means to produce collective dynamics and synchronization of many cells over large distances. We suggest that this is a way to stimulate all the cells slowly so that they polarize cooperatively on the entire spheroid. We believe the onset of this process is based on critical fluctuations rather than deterministic molecular signatures, which may well resolve apparent contradictions in the literature on the nature of axis determining stimuli.

## 5 Experimental procedures

### *Hydra* strains

The wildtype strain of *Hydra vulgaris* and the transgenic strains *Hydra magnipapillata* ks1 GFP, *Hydra vulgaris* 12ASAktin+GFP Ecto- and *Hydra vulgaris* 12ASAktin+GFP endodermal (all kindly provided by the Bosch group, Christian Albrechts University, Kiel, Germany) were used for experiments and imaging. Unless otherwise stated, the transgenic *Hydra magnipapillata* ks1-GFP were used for the experiments.

### Preparation and culturing of *hydra*

Unless otherwise stated, all experiments with *hydra* were performed in Volvic mineral water (Danone Waters, Deutschland GmbH). *Hydra* were fed with artemia, but starved 24h prior to experiments. To obtain tissue pieces, a doughnut-like slice was cut from the centre of the body column while observing through a stereoscope (Stemi 2000, Zeiss, Oberkochen, Germany). This slice was then cut into eight equal pieces, which resulted in fragments that form spheroids of 0.18mm to 0.30mm in diameter. The technique follows the procedure described in ^9,10,30^.

### Staining

Microtubules: *Hydra* balls were relaxed for 1min with 2% urethane in *hydra* medium, and fixed for 1h at room temperature with Landowsky fixative (48% ethanol, 3.6% formaldehyde, 3.8% acetic acid, in water). After three washes with phosphate-buffered saline (PBS), they were permeabilized with 0.8% Triton X-100 in PBS for 15min, blocked with 0.1% Triton X-100, 1% BSA (w/v) in PBS for 20min, and incubated overnight with a monoclonal anti-tubulin antibody (clone 2-28-33; Sigma) at 4°C. Subsequently, these were washed three times with PBS and incubated for 2h with the anti-mouse IgG (whole molecule)-FITC secondary antibody (Sigma). Myonemes: Labelling of the myonemes was performed using rhodamine/ phalloidin, following the protocol above for the permeabilization.

### Micropipette aspiration

Micropipettes were pulled from borosilicate glass capillaries (outer diameter of 1mm and an inner diameter of 0.5mm) using a pipette puller (model P-97; Sutter Instruments). After pulling, the pipettes were trimmed to achieve blunt ends with an outer diameter of 200*µ*m. For the *hydra* to tolerate the contact, the tips needed to be smoothed. A microforge was used for smoothing. For experiments, the pipettes were connected to a homemade low-pressure system using a syringe and a U-shaped tube with marks used as a manometer to control the pressure. The set-up was mounted onto a microscope (IX70; Olympus). To account for the sensitivity of the *hydra* to contaminants and to avoid evaporation, the experiments were performed in closed glass Petri dishes with a hole in the lid for the micropipette. Putting the hydra in place was performed as follows: the *hydra* tissue was allowed to rest and heal for 30 min after cutting. The pipette tip was moved in close proximity to the *hydra* and held within focus using a micromanipulator (Xeno Works; Sutter Instruments). After contact, the pressure within the pipette was quickly increased to avoid escape. The length of the aspirated tissue within the pipette was set to about 60 micrometers, which corresponded to roughly half the radius of the hydra spheroid. The pressure was decreased to the lowest pressure necessary to secure the *hydra* in place, corresponding to a water column roughly a cm high, depending on the precise geometry of the pipette. If necessary, adjustments in the positioning of the *hydra* were made to move it back into focus using the micromanipulator. Large deviations from the aspiration pressure, the micropipette tip shape or diameter lead to either escape or death of the regenerates. During regeneration, oscillations occurred and the different phases of regeneration (strong inflations, weaker inflations, tentacle formation) could be distinguished. No differences from wild type were spotted by eye. The tissue within the pipette contained no lumen, and there were no osmotic inflations in this part. The time until symmetry breaking or the occurrence of tentacles corresponded to the control experiments (about 18 h until the emergence of an axis). The success rate of regeneration until tentacle formation was 60 % (dead and escaped regenerates were counted as not successful). The angle of the axis in the focal plane was plotted (figure). 3 hydras developed in direction of the light path, head up, the others developed in different directions (N=9), but within a circular area around the pipette tip (see figure, supplementary). No hydra regenerated with the tentacles facing downwards, pointing either to a sensitivity to gravity, as suggested by H. Shimitzu ^29^, or a sensitivity to the red light from a light emitting diode used for illumination. Controls were left to regenerate individually on the bottom of a petri dish, slightly below the location given by the pipette tip. Here heads developed exclusively in directions parallel to the bottom surface, suggesting a sensitivity to the presence of the surface. Turning hydras were excluded from the analysis. The angular orientation was determined and binned in directions of 30°C for plotting the distribution (N= 17).

### Microscopy

Confocal microscopy of hydra spheroids was performed in 35mm diameter dishes (Ibidi *µ*). Visualization of the actin structures was performed using an inverted microscope (TI-Eclipse; Nikon), with a spinning head (Yokogawa CSU-W1; Andor), a combiner (LU-NV; Nikon), 70 mW, and a 4-million-pixel digital camera (ORCA Flash 4.0 V2 CMOS; Hamamatsu) controlled by NIS Elements software. For imaging actin myonemes, a 20x CFIPlan Apo NA 0.75 objective with a working distance of 1.0 mm was used. The excitation/ emission wavelengths were 540/565 nm. A 60x oil immersion objective (Plan Apo; NA 1.4) was used to collect detailed images of actin and the images of the microtubules under excitation/emission wavelengths of 488/540 nm. Here the bottom of the petri dish needed to be replaced with a cover glass. The microscopy of the hydra spheroid regeneration patterns was performed using a phase contrast microscope (IX70; Olympus) equipped with a digital camera (G1-2000; Moravian Instruments). For the regeneration pattern analysis, the microscopy images were recorded at a frame rate of 1/s. Imaging of the spheroids was performed with a PLN 10x or 4x objective (Olympus).

### Hanging drop technique

For regeneration, the samples were prepared in culture medium, which was supplemented with drugs in some experiments. Drops of 30*µ*l that included one spheroid each were pipetted onto the lid of a plastic petri dish (diameter, 5cm; Corning). The lids were put on top of medium-containing dishes, which were then sealed with parafilm to avoid evaporation. The *hydra* floated in the drops hanging down from the lids. Hanging drops allowed the fluctuations of the main axis orientation to be seen more clearly compared to the bottom of a petri dish. The drop kept the tissue fragments in focus. Control experiments were performed using silica-beads.

### Image analysis

A series of images was transformed to binary, and particle analysis was performed using open-source FIJI Imaging software. Area, circularity, axis length and main axis orientation were determined. The data obtained this way were analyzed and plotted using Origin Pro 8.6 (Origin Lab) and MatLab R2011b (Mathworks). Confocal images were processed using FIJI. Often the image treatment was precluded by ejected debris that remained stuck to the spheroid.

All of the correlation functions were calculated using Origin Pro 8.6. To analyze the correlations between volume and angle, and others, cross-correlation was performed over the regeneration from the beginning until the appearance of tentacles. Cross-correlation provides a measure of similarity between two series as a function of the lag of one series relative to the other. The following function was used for the discrete functions *a* and *b*:

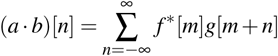

where *f*\* is the complex conjugate, and *n* is the lag. The correlations between the size changes of the spheroids and time span until reaching final size were assessed using Pearsons correlation coefficients. This coefficient represents a measure of the strength and direction of the linear relationship between two variables, and it is defined as the covariance of the variables divided by the product of their standard deviations. The correlations between the size changes and the time until mechanical symmetry breaking were performed using Spearman correlation coefficients. This coefficient represents a measure of the strength and direction of the monotonic relationship between two variables.

In case of actin modifying drugs the analysis was performed on the first debris-free recording, checking however, that the regeneration did not deviate in any obvious manner from the other observed regenerations by eye. Since the conclusions regarding actin were limited to observing that axis formation was not prevented and that the timing of axis formation exhibited a huge change, no further analysis was performed regarding these drugs.

Regarding the effects of microtubule modifying drugs, a statistical analysis was performed to describe the patterns. To describe the fluctuating behaviour of the spheroids with paclitaxel, the differences between two subsequent values of the angular position were determined. To avoid artificially high angular differences that can result from angles that change from 180 degrees to 0 degrees (or vice versa), cyclic boundaries were applied. The mean scattering of the angle during the stable phases was determined and used as threshold. If the angular differences were within this threshold, the spheroid was considered to be stable. The periods of time the spheroids remained in either stable or unstable states were determined and plotted against the frequency of appearance of a period of different lengths. Nocodazole reduced the fluctuations and the above analysis was not suitable. Accordingly a spectral analysis of the inflation was performed by subtracting a linear fit from each individual inflation, detrending the data. Fast Fourier transform was performed to compute the spectra. Respecting the Nyquist-Shannon theorem, frequencies above 0.1 Hz were discarded.

### Drugs

The drugs for the analysis depicted in figures 5 and 6 were used at the highest viable concentrations as determined on adult *hydra*, prior to the experiments.

Nocodazole (Sigma Aldrich) was used at a working concentration of 10^−6^ M, and stored at 6 × 10^−3^ M in dimethylsulphoxide (DMSO). Paclitaxel (Sigma Aldrich) was used at a working concentration of 10^−6^ M, and stored as a 10^−3^ M solution in DMSO. Y27632 was used at a working concentration of 10^−8^M from stock at 10^−3^ M in DMSO. SMIFH2 was used at a working concentration of 2.5·10^−5^ M, stored in DMSO at 10^−3^ M. Phalloidin was used at a working concentration of 2 · 10^−5^ M.

For experiments where the action of nocodazole was probed alongside paclitaxel, the *hydra* balls (200 − 300*µ*m) were treated with a combination of nocodazole (1*n*M, 0.1*n*M, 0.01*nM*) and paclitaxel (1*µ*M, 0.1*µ*M, 0.01*µ*M) at different concentrations (*n* = 30). The minimal effective concentrations of nocodazole for inhibition of the regeneration was determined as 0.1*n*M (*n* = 15, out of 15). To suppress the effects of nocodazole, the minimum concentration of paclitaxel was determined as 0.1*µ*M, when *hydra* balls (*n* = 15) were first incubated with nocodazole (0.1*µ*M) for 24h, and then with paclitaxel (0.1*µ*M). We observed that if paclitaxel rescued the process, the time to regenerate *hydra* was prolonged significantly (by about 40h) in nocodazole-supplemented medium, compared to (untreated) controls). However, the paclitaxel treated hydra spheroids survived and regenerated (*n* = 14, out of 15).

## 6 Author Contributions

H.D. and A.O conceived of the study and interpreted the data. Regeneration experiments and analysis were performed by H.D., A.P. and V.G.. Rhodamine-phalloidin staining was performed by M.S. and H.D. Microtubule staining was performed by A.P. Confocal microscopy was performed by E.T.. Analytical guidance was provided by M.S, V.G. and A.O.. The manuscript was drafted by A.O. and H.D.. F.L. and A.O. supervised.

## 7 Acknowledgements

We acknowledge fruitful discussions with Heiko Rieger in connection with the interpretation of the spectral analysis data. Funding was provided by Universität des Saarlandes and Deutsche Forschungsgemeinschaft (DFG) in the framework of the Collaborative Research Centre (SFB 1027). All *hydra*, including the transgenic ones, were kindly provided by Thomas Bosch and his group from the Christian-Albrechts Universität zu Kiel, Germany.

