## Supplementary Material for "Symmetry breaking and *de-novo* axis formation in *hydra* spheroids: the microtubule cytoskeleton as a pivotal element"

Supplementary Material (ESI)

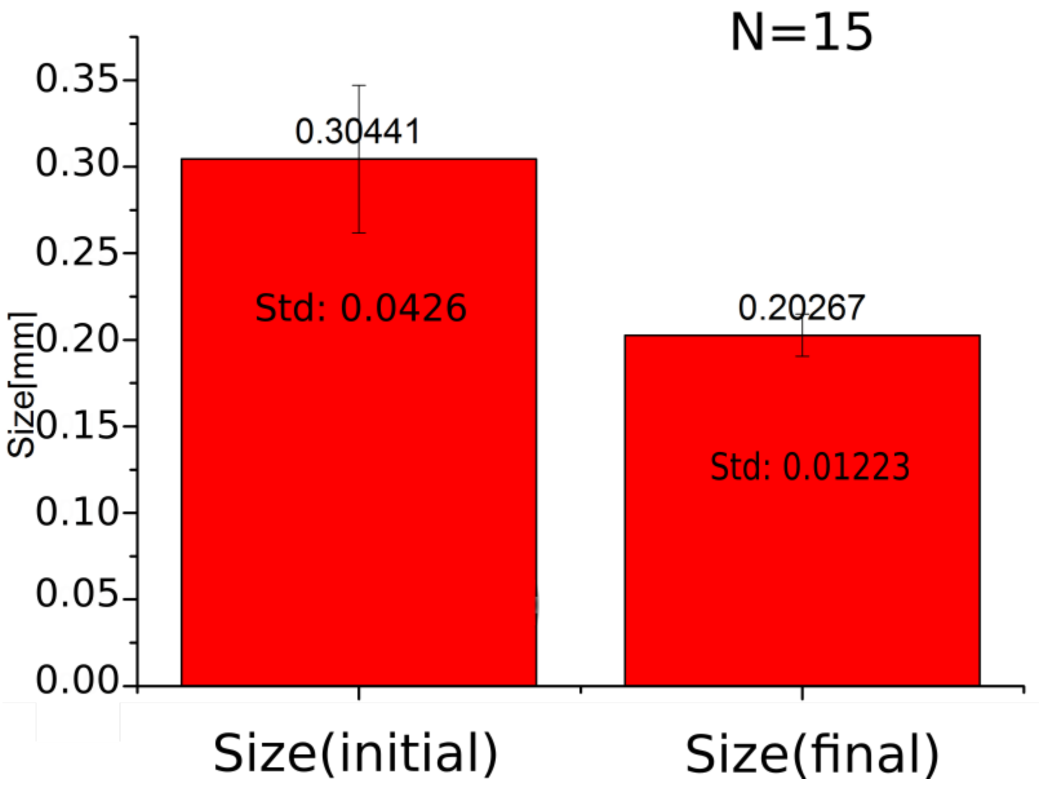


Figure 1: Comparison of the diameter of the hydra spheroids after the first deflation (left) compared to the deflated diameter of the hydra spheroid before the axis appears (right). The standard deviation is lower on the right, indicating that different sized spheroids tend to adjust to the same diameter N=15 spheroids.

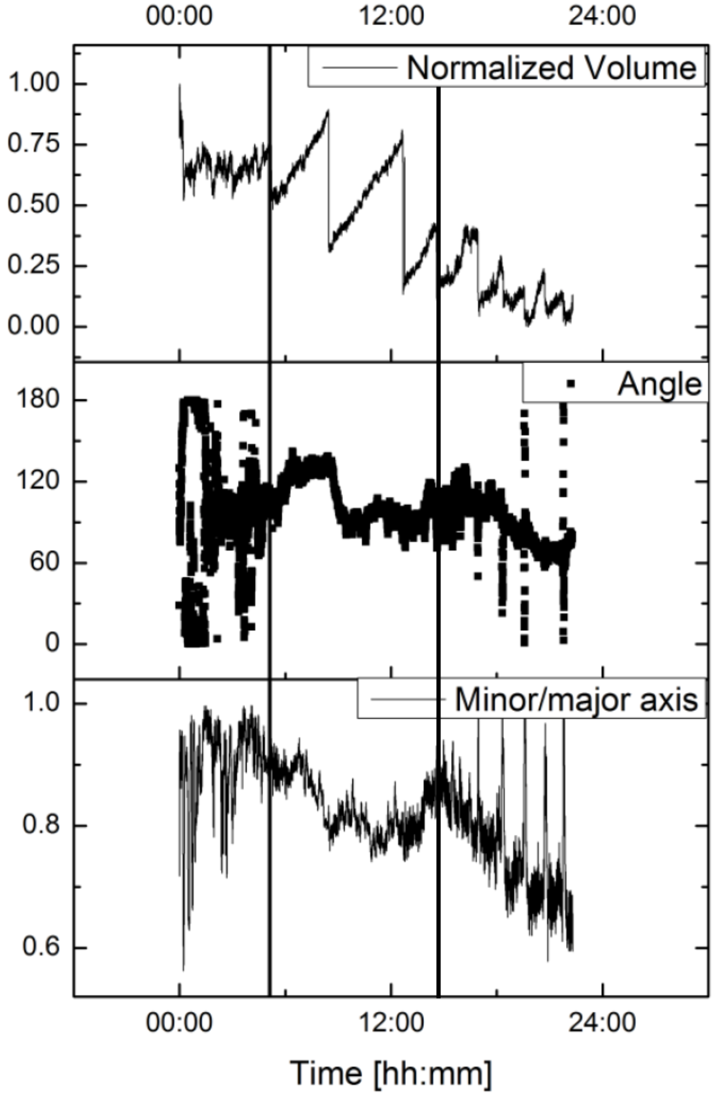


Figure 2: Successful axis definition of a spheroid that did not follow the usual inflation pattern. It appears that an initial week spot from the preparation is lost before the inflation follows its usual pattern (large volume inflations transiting to smaller inflations) and the axis is redefined (black vertical lines). A possible interpretation is that an early organizer is lost before it is recreated, which may well explain the accelerated formation of the organize compared to control (18h).

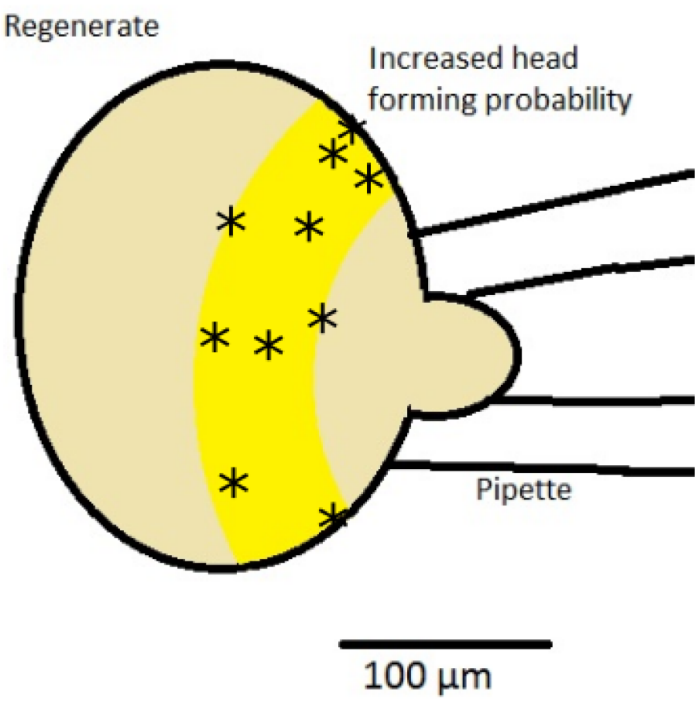


Figure 3: Locations of the emerging hydra heads on the regenerating speroid held by a micropipette. Each cross corresponds to a single head, except for the central cross that corresponds to vertical orientation: here, three heads emerged.

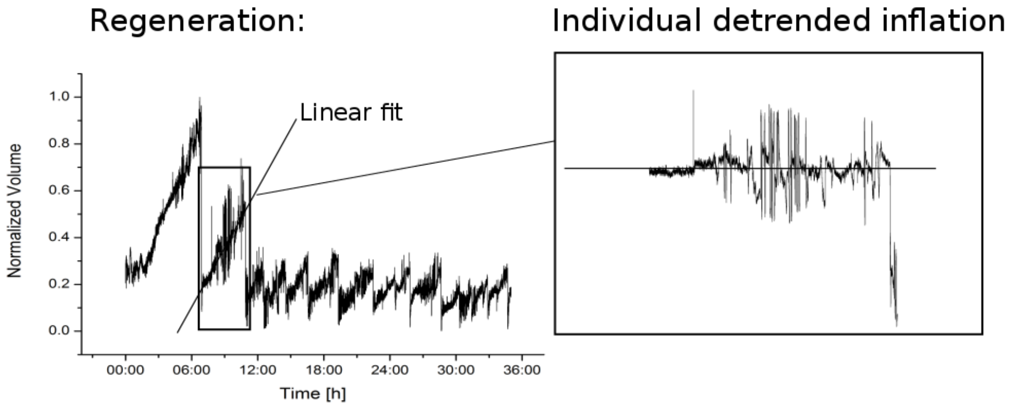


Figure 4: Spectral analysis and cross correlation. A linear fit on every individual inflation is subtracted from the data to obtain a detrended signal. The spectral analysis is performed on the detrended signal (for each inflation).

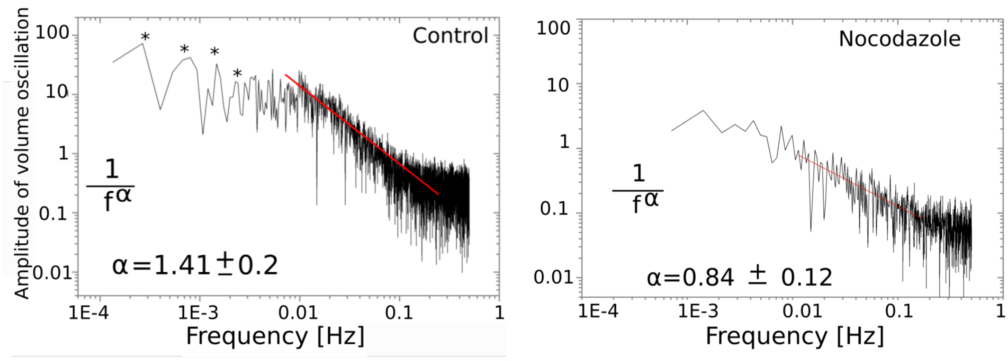


Figure 5: For Fourier transform, each sawtooth of the volume oscillations of the spheroids was detrended by subtraction of a linear fit (Supplementary Information figure S4). The analysis of the detrended oscillations revealed $1/f$ scaling behaviour of the frequency spectrum, $1/f^{\alpha}$, where $\alpha = 1.43 \pm 0.21$ ($n = 8$) for the controls, above frequencies of $0.01Hz$. The control spectrum showed individually distinguishable peaks at the lower frequencies (marked with asterisks). Multiplying this frequency by the duration of the observed periods of inflation revealed the number of sinusoids during each inflation. Each inflation included one sinusoid oscillation that was superimposed by 2, 3, 4 and 6 higher harmonics. This means that each inflation followed a regular pattern of superimposed sinusoids that scaled with the absolute duration of the inflation (and was hence independent of the size of the spheroid). Upon addition of nocodazole, these individual peaks disappeared, and the amplitude was reduced by a factor of 10, although $1/f$ scaling prevailed. The mean exponent in *hydra* treated with nocodazole was $0.84 \pm 0.12$ ($n = 9$ regenerations).

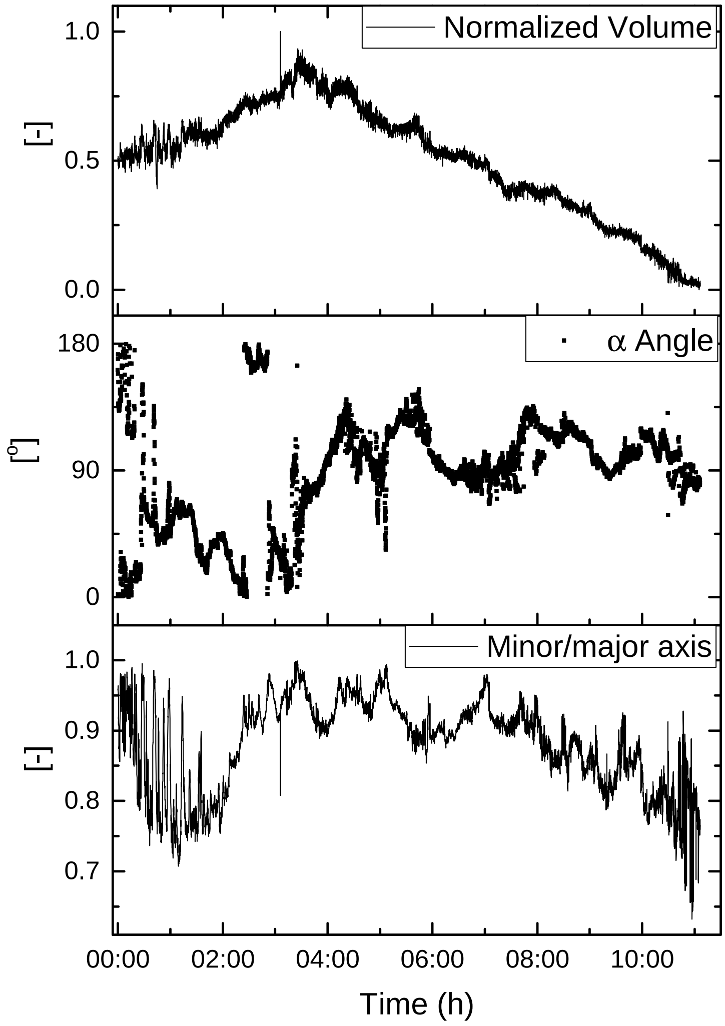


Figure 6: Shape development of a nocodazole (0.1 nM) treated spheroid in nocodazole free medium as a function of time where (00:00) corresponds to the beginning of recording, that is, 24h after application of nocodazole directly after cutting the fragment. There is no sign of axis development. The spheroid slowly decays as indicated by the shrinking volume. The regeneration can be rescued by the application of paclitaxel at time 00:00 (see main, figure 7.
